# Introducing a proline in the α1 M2-M3 linker relieves a molecular brake on channel activation in α1β2γ2 GABA_A_ receptors

**DOI:** 10.64898/2026.03.10.710710

**Authors:** Netrang G. Desai, Pranavi Garlapati, Cecilia M. Borghese, Marcel P. Goldschen-Ohm

## Abstract

GABA_A_ receptors (GABA_A_Rs) are pentameric ligand-gated ion channels (pLGICs) essential for inhibitory synaptic transmission throughout the central nervous system. Despite progress in understanding their three-dimensional structure, the molecular basis for how neurotransmitter binding is transduced to ion channel gating remains poorly understood. Furthermore, relatively little is known about the contributions of distinct subunits to this coupling within typical heteromeric receptors. A highly conserved proline (site 1) in the M2-M3 linker of pLGIC subunits is involved in channel gating – e.g., P273 in the GABA_A_R β2 subunit. In GABA_A_Rs, only the β subunits have an additional proline in the M2-M3 linker (site 2) – e.g., β2(P276) – whereas all other subunits have a non-proline at the homologous site 2 position. Here, we investigate the functional contribution of proline at site 2 in distinct subunits of α1β2γ2 GABA_A_Rs. We expressed wild type or mutant α1β2γ2 GABA_A_Rs in *Xenopus laevis* oocytes and used two-electrode voltage clamp electrophysiology to record channel currents in response to GABA and/or other ligands. First, we introduced a proline at site 2 in α1 or γ2 subunits: α1(A280P) and γ2(S291P). Second, we replaced the site 2 proline in the β2 subunit with its homologous non-proline residue from α1 or γ2 subunits: β2(P276A) or β2(P276S). We show that α1(A280P) confers enhanced GABA-sensitivity and spontaneous unliganded channel activity, whereas γ2(S291P) has minor effects on channel activation. In contrast, β2(P276A) or β2(P276S) either had no effect or enhanced GABA-activation, respectively, indicating complex functional dependence on the side chain at site 2 in the β2 subunit. When in combination with other substitutions, the presence or absence of α1(A280P) was consistently correlated with enhanced GABA-sensitivity and spontaneous activity. Thus, introduction of a proline at site 2 in the α1 M2-M3 linker biases the channel towards an activated state and prevents it from remaining closed at rest.

## Introduction

For GABA_A_ receptors (GABA_A_Rs) and related pentameric ligand-gated ion channels (pLGICs), coupling of neurotransmitter binding in the extracellular domain (ECD) to gating of the ion channel pore in the transmembrane domain (TMD) involves several loops located at the ECD-TMD interface (**Figure 1A**) (Kash et al., 2003; Kash, Dizon, et al., 2004; Kash, Trudell, et al., 2004; Moroni et al., 2011; Venkatachalan & Czajkowski, 2012). One of these loops, the M2-M3 linker, connects the second and third transmembrane domains, and interacts with ECD loops (**Figure 1B**) (Borghese et al., 2025; Fisher, 2002; Hales et al., 2006; Kash et al., 2003; Kash, Trudell, et al., 2004; Nors et al., 2021, 2024; O’Shea & Harrison, 2000; O’Shea et al., 2009; Xiu et al., 2005). Studies have uncovered asymmetric functional effects of M2-M3 linker perturbations in distinct subunits within typical heteromeric synaptic α1β2γ2 GABA_A_Rs (Borghese et al., 2025; Hales et al., 2006; Nors et al., 2021, 2024). However, the physical basis underlying this asymmetry remains to be fully understood.

**Figure 1.**
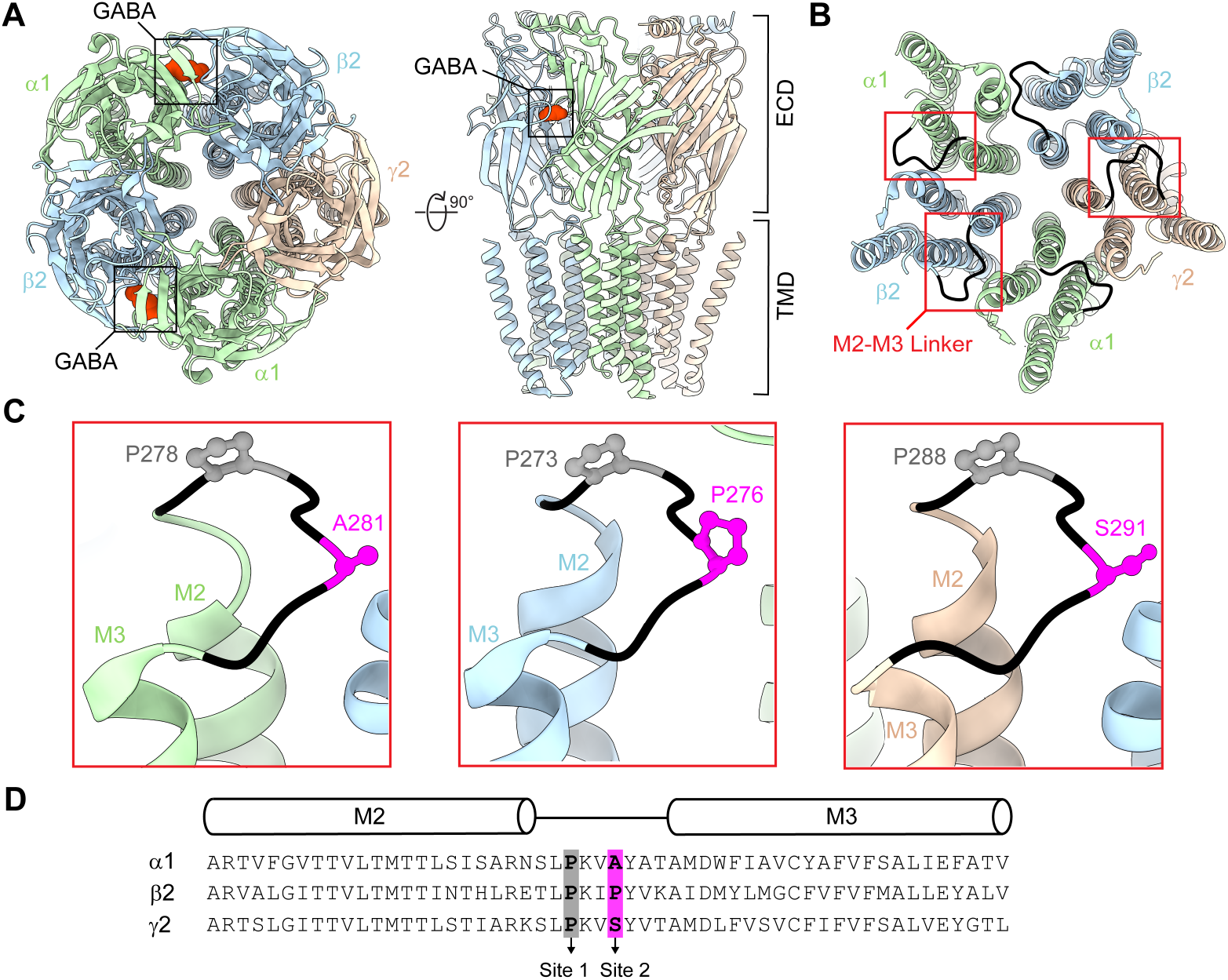
M2-M3 linker prolines in a synaptic α1β2γ2 GABA_A_R. **(A)** Cryo-EM map (PDB: 6X3X) of a human synaptic α1β2γ2 GABA_A_R in complex with GABA (red) viewed from the top (left) and side (right). (**B**) Same view as in panel A for the transmembrane domain only. The M2-M3 linkers are colored black. **(C)** Zoom in of boxed regions in panel B. A highly conserved proline at site 1 in α1, β2, and γ2 subunits is highlighted in grey. A proline at site 2 found in β subunits only and the homologous non-proline residues in α1 and γ2 subunits are highlighted in magenta. The mature protein numeration for β2 and γ2 subunits is the same for rat and human, but the numeration for the rat α1 subunit is (human numeration -1) for most of the subunit. **(D)** Sequence alignment of the M2-M3 linker regions (black line) for rat α1, β2, and γ2 subunits. Transmembrane helices M2 and M3 are indicated with cylinders above the alignment. Structures visualized using ChimeraX (Pettersen et al., 2021).

All pLGICs have a highly conserved proline in the M2-M3 linker at site 1 which is involved in channel gating (**Figure 1C-D, 2**). Early studies of nicotinic acetylcholine receptors (nAChRs) suggested that a “pin-in-socket” mechanism involving this conserved proline contributes to coupling agonist binding to pore opening (Hanek, 2009; Hanek et al., 2008; Lee et al., 2008; Lee & Sine, 2005; Lee et al., 2009; Miyazawa et al., 1999, 2003; Nigel Unwin, 2005; N Unwin et al., 2002). Consistent with this idea, substitutions of this proline in homomeric α7 nAChR result in non-functional channels (Mosesso et al., 2019). However, substitutions in other pLGICs such as GABA_A_Rs and 5-HT_3A_ receptors do not abolish channel function, indicating that such a mechanism is not requisite in these channels (Brodzki & Mozrzymas, 2022; Kaczor et al., 2022; Mosesso et al., 2019; Xiu et al., 2005). Nonetheless, substitutions of this conserved proline in either α1 or β2 subunits of GABA_A_Rs do alter channel gating properties (Brodzki & Mozrzymas, 2022; Kaczor et al., 2022), indicating that this proline at least contributes to channel activity.

In addition to the highly conserved site 1 proline, cationic nAChRs and 5-HT_3A_ receptors have another proline located at the top of the M3 transmembrane helix (site 3) (**Figure 2**). One of the proposed mechanisms for gating is the cis-trans isomerization of this site 3 proline which strongly correlates with the channel activation of 5-HT_3A_ receptors (Lummis et al., 2005). Although GABA_A_Rs lack a homologous proline at site 3, they do have an additional proline in the M2-M3 linker which is specific to β subunits only (site 2) (**Figure 1C-D, 2**). Here, we investigate the functional contribution of proline at site 2 in each distinct subunit within typical synaptic α1β2γ2 GABA_A_Rs. We show that introduction of a proline at site 2 in the α1 subunit enhances GABA-sensitivity and prevents the channel from remaining closed at rest. The role of proline at site 2 in the β2 subunit is more ambiguous, with differential effects for alanine or serine substitutions. Nonetheless, across tested combinations of substitutions in multiple subunits, enhanced GABA-sensitivity and spontaneous activity consistently correlated with the presence of a proline at site 2 in the α1 subunit. This suggests that the presence or absence of a proline-induced kink in the α1 M2-M3 linker at site 2 is an important determinant of channel activation. Our observations provide new insight into the subunit-specific effects of M2-M3 linker perturbations on the channel gating of α1β2γ2 GABA_A_Rs.

**Figure 2.**
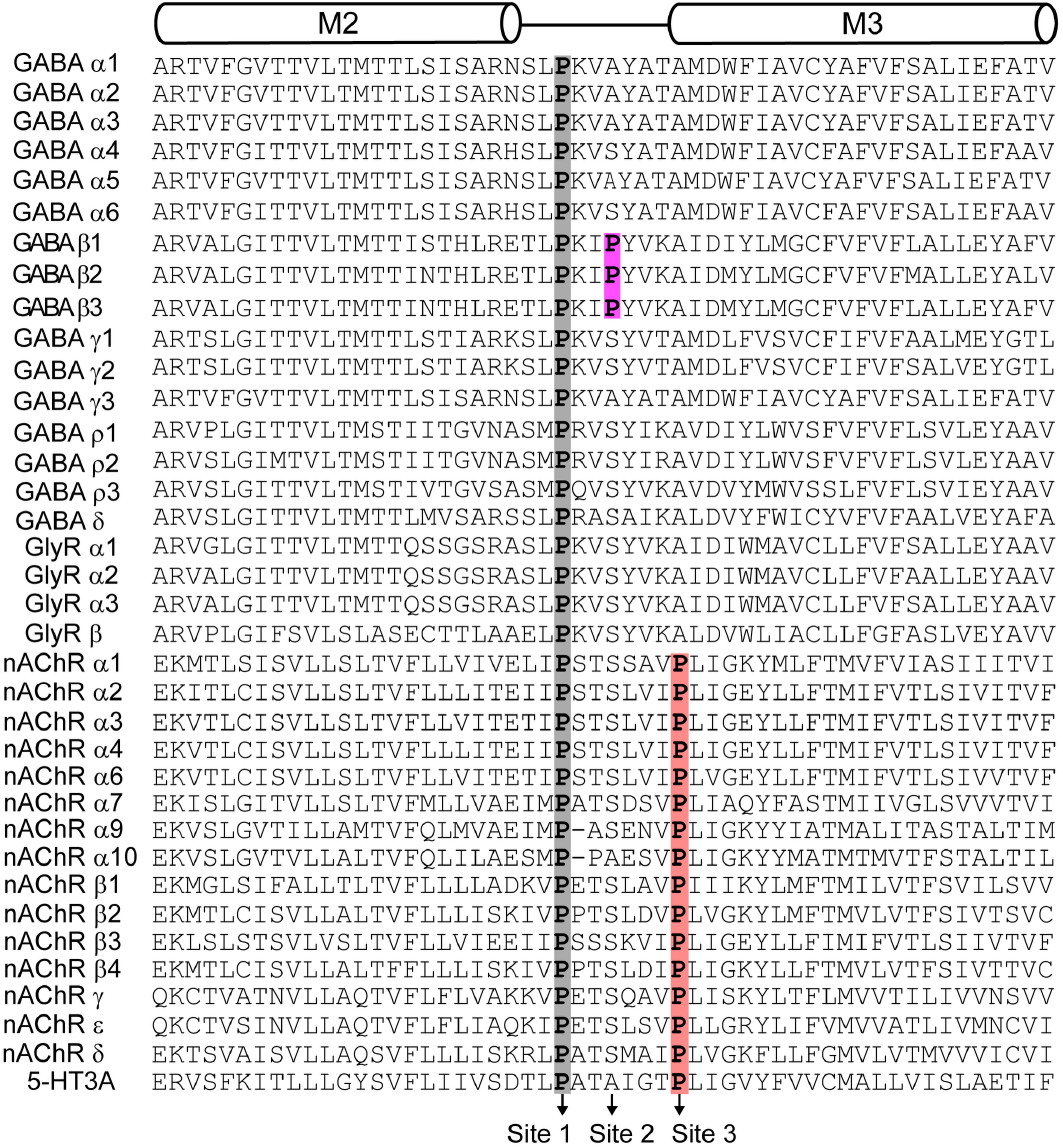
Sequence alignment of M2-M3 transmembrane region for pLGICs. Transmembrane helices M2 and M3 are indicated with cylinders above the alignment. A conserved proline at site 1 across the subunits of pLGIC receptors is highlighted in grey. A proline at site 2 conserved only in β subunits of GABA_A_Rs is highlighted in magenta. A proline found at site 3 in nAChR and 5-HT_3A_ subunits is highlighted in salmon. All sequences are from rat, which are very similar to those from humans. Importantly, the overall site 1-3 proline pattern is the same in human subunits.

## Results

### Introducing a proline at site 2 in the α1 M2-M3 linker enhances channel activation

*Xenopus laevis* oocytes were coinjected with cRNA for α1, β2, and γ2 GABA_A_R subunits (either wild type or mutants) in a 1:1:10 ratio (Boileau et al., 2002), and current responses to application of ligands were recorded using two-electrode voltage clamp (see representative current tracings in Figure 3). For each oocyte, we measured current responses to 20-40 s pulses of a series of concentrations of GABA to assess activation sensitivity, and finally the response to the combination of saturating GABA and propofol to estimate the maximum current amplitude when all channels are open with ∼100% probability as described previously (Pierce et al., 2024) (**Figure 4A**). GABA concentration-response curves (CRCs) were normalized to the peak current response elicited by co-application of saturating GABA and propofol, and then fit with the Hill equation (Eq. 1) (**Figure 4B-C**). Relative comparison of the maximal response to either GABA alone or GABA and propofol allows estimation of the maximum efficiency with which GABA can open the channel.

**Figure 3.**
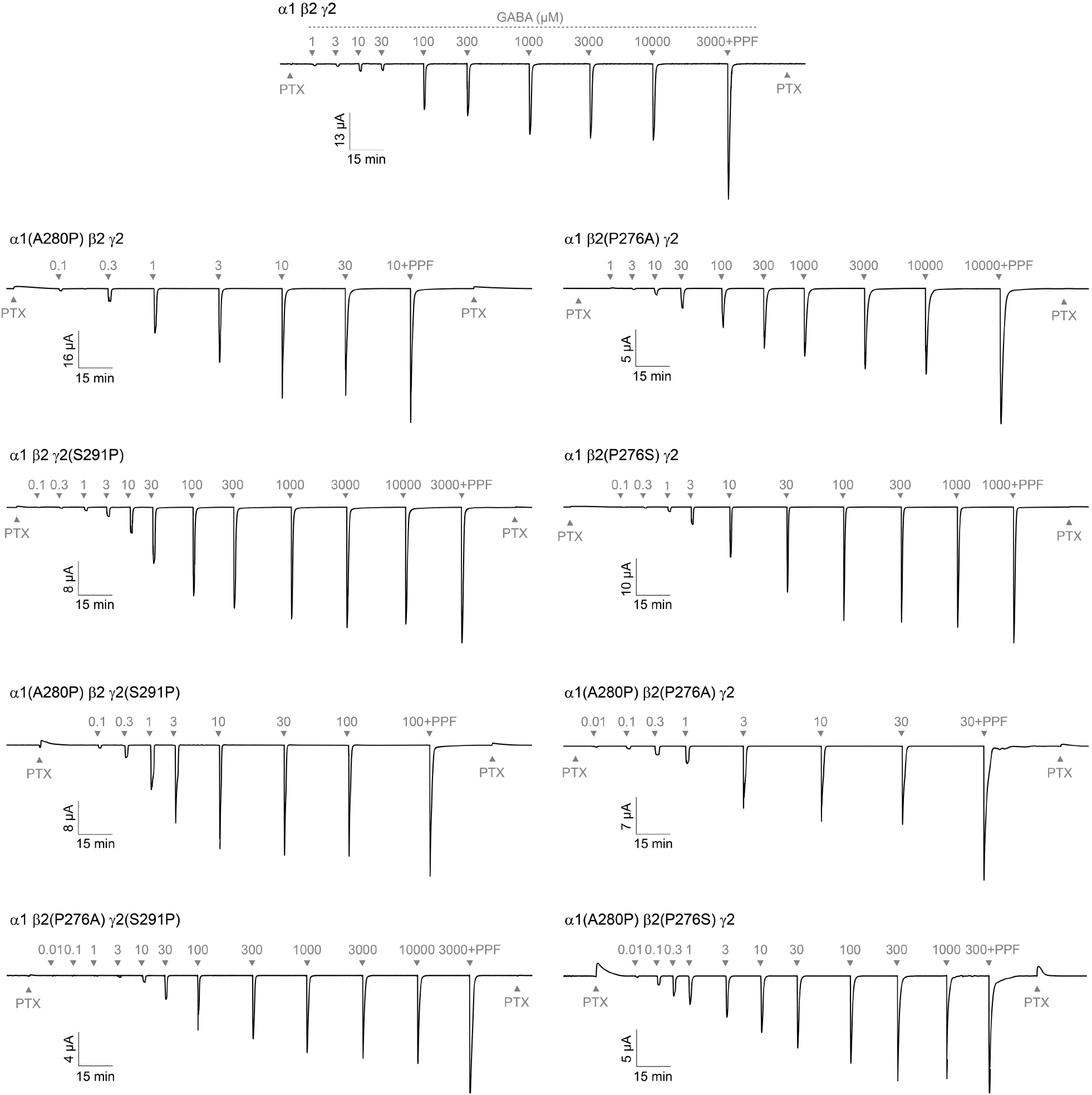
Representative current tracings of wild-type and mutant α1β2γ2 GABA_A_Rs. Example two-electrode voltage clamp recordings of wild-type and mutant α1β2γ2 GABA_A_Rs. A 10 s pulse of 1 mM picrotoxin (PTX; to assess for unliganded opening) was followed by a series of 20-40 s pulses (sufficient to resolve peak) of increasing concentrations of GABA (µM), a 20 s pulse of saturating GABA in combination with 30 µM propofol (PPF) to estimate the maximal possible current, and a final 10 s pulse of 1 mM PTX. Labeled numeric values for pulses indicate GABA concentration (µM).

**Figure 4.**
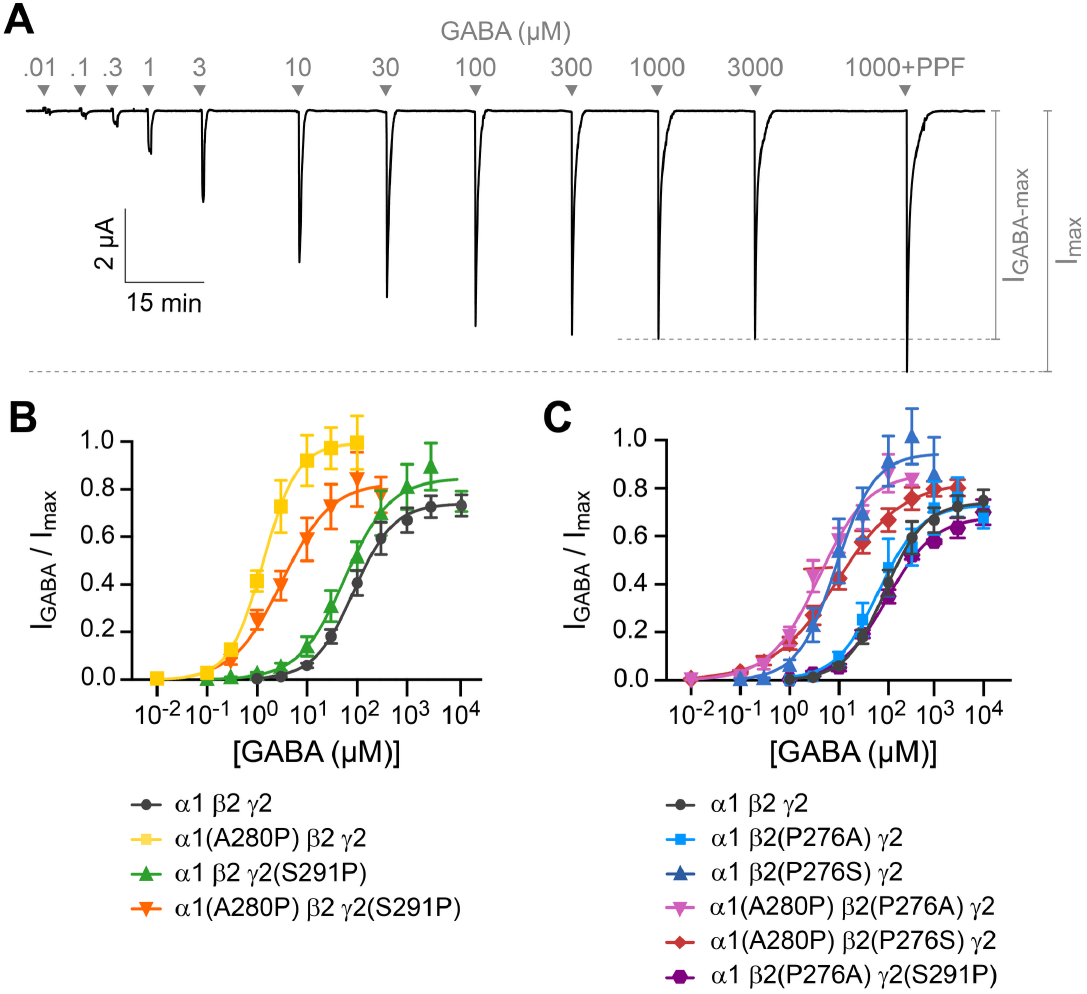
The substitution α1(A280P) promotes channel activation. **(A)** Representative current tracing of the mutant α1(A280P) β2(P276S) γ2 GABA_A_R construct. Peak current responses to increasing concentrations of GABA (µM) followed by a pulse of saturating GABA in combination with 30 µM propofol (PPF). **(B, C)** Normalized concentration-response curves for GABA-elicited currents. Curves are the Hill equation fit to the means (see Eq. 1). Datapoints are plotted as mean ± SEM. See Table 1 for summary statistics of fit parameters and number of oocytes.

First, we investigated the effects of introducing a proline at site 2 in the α1 and/or γ2 subunit M2-M3 linker analogous to the naturally occurring proline at the homologous position in the β2 subunit. The substitution α1(A280P) increases GABA sensitivity (i.e., left-shifts the GABA CRC) by ~70-fold (**Figure 4B**). Furthermore, whereas GABA opens wild type channels with a peak open probability ∼0.7, α1(A280P) enhances the efficiency of GABA-activation such that current responses reach a peak open probability of ∼1.0 (**Figure 4B**). In contrast, γ2(S291P) has little effect on channel activation, although there was a small trend towards enhanced activation. The double substitution α1(A280P)γ2(S291P) increases GABA sensitivity by ~30-fold, with a similarly modest increase to maximal opening as for γ2(S291P) alone (**Figure 4B**). Thus, there is some non-additivity in the effects of the double mutant, although α1(A280P) is the more dominant substitution overall. In summary, introduction of a proline at site 2 in the α1 subunit sensitizes the channel for activation by GABA.

**Table 1.**
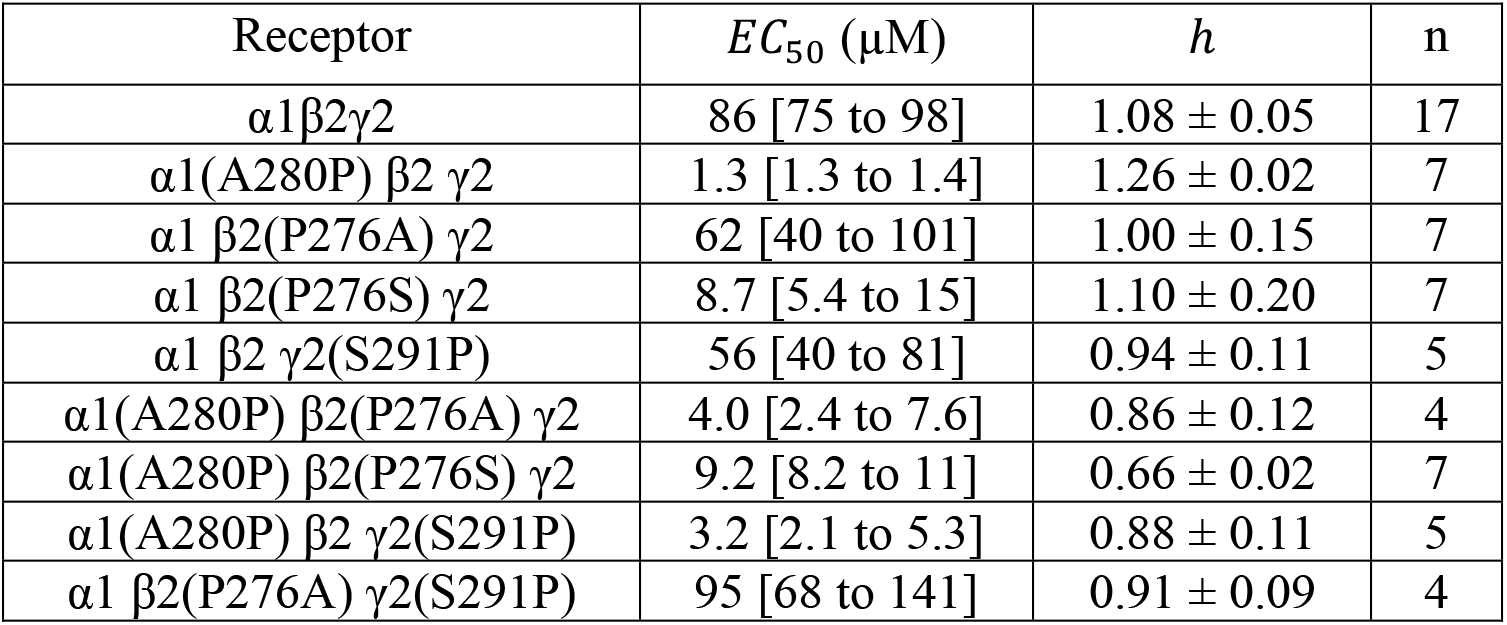
Parameters for Hill equation fits to normalized GABA CRCs from Figure 4. Hill equation is defined in Eq. 1. *EC*_50_ is expressed as mean [95% confidence interval], *h* as mean ± SEM, and n is the number of oocytes.

### Substituting the site 2 proline in the β2 M2-M3 linker has sidechain-dependent effects on channel activation

We investigated the effects of replacing the site 2 proline in the β2 subunit with the homologous alanine or serine in α1 or γ2 subunits, respectively. β2(P276S) enhances GABA-sensitivity (i.e., left-shifts the CRC) by ~10-fold and confers an enhanced maximal GABA-elicited peak open probability of ∼0.9 (**Figure 4C**). In contrast, β2(P276A) has no effect on channel activation. Thus, there is a more complex dependence of channel activation on the properties of the sidechain at site 2 in β2 subunits than simply whether it is a proline. We further evaluated double substitutions that effectively move the proline from β2 to α1 or γ2 subunits. Both α1(A280P)β2(P276S) and α1(A280P)β2(P276A) confer increased GABA-sensitivity and enhanced maximal open probability similar to that of α1(A280P) or β2(P276S) alone (**Figure 4C**). Although the underlying mechanism for the differential effects of alanine or serine substitution at β2(P276) are unclear, the observation that introduction of a proline at site 2 in the α1 subunit enhances channel activation is still robustly supported by the tested double substitutions.

### Introducing a proline at site 2 in the α1 M2-M3 linker prevents channels from remaining closed at rest

To assay for any spontaneous unliganded channel opening in the mutants, we applied the pore blocker picrotoxin (PTX) (**Figure 5A**). The unliganded channel open probability (P_open_) was estimated as the ratio of PTX-sensitive current to the total current from the baseline in PTX to the peak current elicited by the combination of saturating GABA and propofol as illustrated in Figure 5A (Borghese et al., 2025; Nors et al., 2024). In comparison to wild-type channels, which are essentially fully closed at rest with an unliganded P_open_ ≤ 0.002 or 0.2% (Borghese et al., 2025; Mortensen et al., 2003), α1(A280P) confers obvious PTX-sensitive spontaneous channel activity (unliganded P_open_ = 0.05 or 5%) (**Figure 5B**). All tested combinations of substitutions that included a proline at site 2 in the α1 subunit show noticeable unliganded openings with probabilities of a few percent. The double substitution α1(A280P) β2(P276S) conferred the most spontaneous channel activity (unliganded P_open_ = 0.06 or 6%) (**Figure 5B**). Although this unliganded activity is rather small in comparison to maximal GABA-evoked opening, it is nonetheless indicative of a relatively sizeable energetic perturbation to the closed-open equilibrium. This is because the energy difference between closed and open states needs to change appreciably before channel opening can even be reliably detected in two-electrode voltage clamp recordings. We estimate that a change in P_open_ from 0.2% to 5% requires approximately 1/3^rd^ of the total energy supplied from GABA binding (see Borghese et al. (2025) for a more detailed description of this estimation). Thus, introduction of a proline at site 2 in the α1 subunit biases the channel approximately 1/3^rd^ of the way along its activation pathway and prevents the channel from remaining closed at rest.

**Figure 5.**
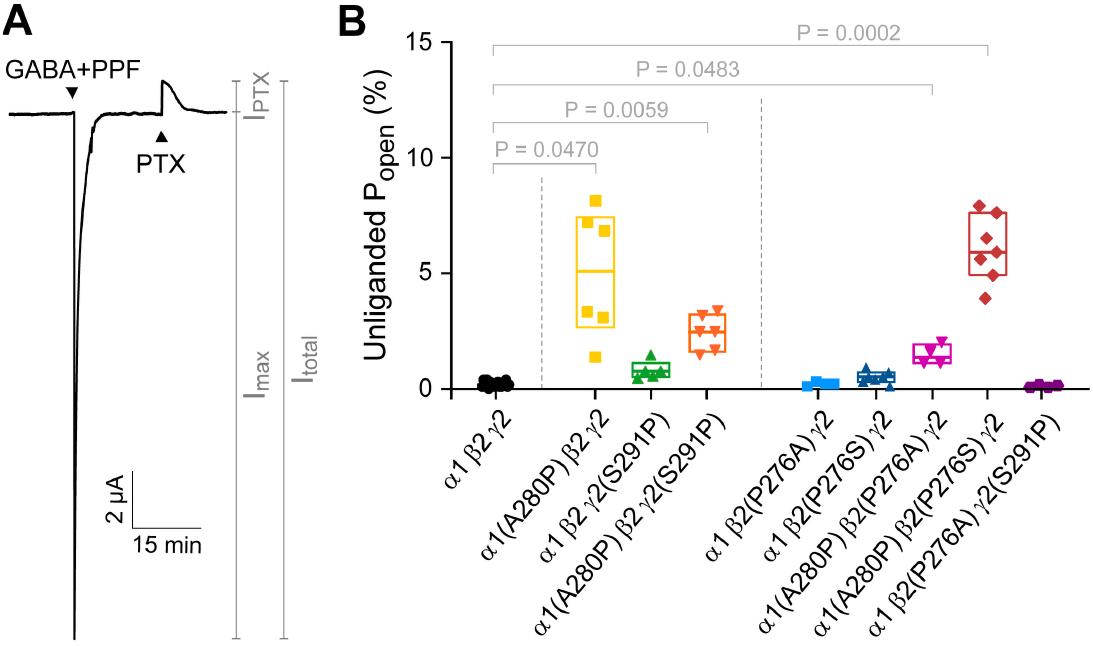
The substitution α1(A280P) confers spontaneously open channels. **(A)** Mutant α1(A280P) β2(P276S) γ2 GABA_A_R construct. The total current (I_total_) is estimated as the sum of the peak current response to saturating GABA and 30 µM propofol (PPF; I_max_) and the current elicited by 1 mM picrotoxin (PTX; I_PTX_). **(B)** The unliganded open probability (P_open_) is estimated as the ratio of PTX-elicited current to the total current (I_PTX_/I_total_). Box plots indicate median and quartiles. *P*-values < 0.05 for Brown-Forsythe ANOVA with posthoc Dunnett’s T3 test are shown. See Table 2 for summary statistics and number of oocytes.

**Table 2.**
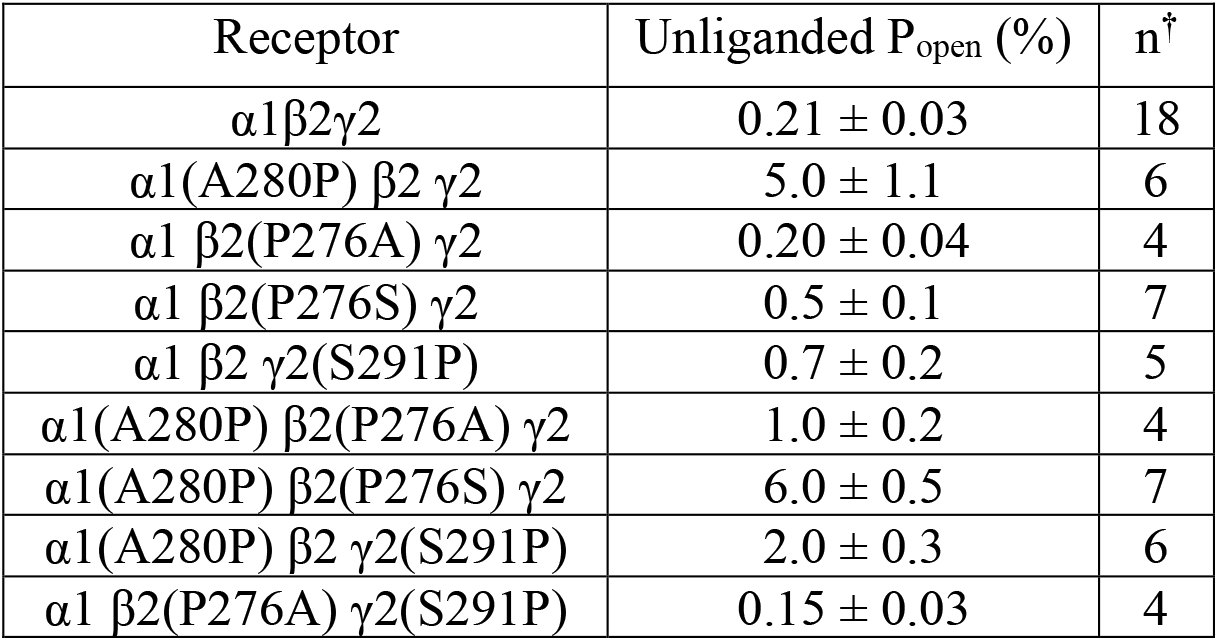
Summary of unliganded open probabilities from Figure 5. Unliganded P_open_ is expressed as mean ± SEM, and n is the number of oocytes. ^†^In a few cases, the oocyte count is different from Table 1 because the unliganded P_open_ could not be estimated due to loss of recording stability during the final application of PTX.

## Discussion

We have previously identified M2-M3 linker substitutions in widely expressed heteromeric α1β2γ2 GABA_A_Rs with asymmetric subunit-specific effects. For example, alanine substitution of the central residue in the M2-M3 linker of β2 or α1 subunits either enhances or inhibits channel activation, respectively (Nors et al., 2024). Ablation of a main-chain hydrogen bond in the β2 subunit M2-M3 linker also enhances channel activation and prevents the channel from remaining closed at rest, whereas analogous hydrogen bond ablation in the α1 subunit has no effect (Borghese et al., 2025). Given the location of the β2 subunit M2-M3 linker directly below the GABA binding sites, we hypothesized that M2-M3 linkers at GABA-binding (β2) and non-binding (α1, γ2) interfaces may have distinct functional contributions to channel gating. One difference in the amino acid sequences of M2-M3 linkers is the site 2 position, which is a proline in β subunits and a non-proline residue in all other subunits. Given that prolines in nearby positions within or next to the M2-M3 linker (sites 1 and 3) have been implicated as important determinants of channel gating (Brodzki & Mozrzymas, 2022; Kaczor et al., 2022; Lummis et al., 2005), we hypothesized that the naturally occurring proline at site 2 in β subunits may contribute to the distinct functional contribution of the β2 M2-M3 linker in α1β2γ2 receptors.

To test this hypothesis, we performed various substitutions which swapped proline and non-proline residues at site 2 in α1, β2, and γ2 subunits. Although our results do not unambiguously define the functional requirements for the site 2 residue in distinct subunits, they do provide new insight into the molecular basis for channel gating in this region. We show that introduction of a proline at site 2 in α1 subunits 1) sensitizes the channel to activation by GABA, 2) maximizes the efficiency of GABA activation, and 3) prevents the channel from remaining closed at rest. We conjecture that introduction of a proline-induced kink allows the α1 M2-M3 linker to more easily move radially outward from a naturally more rigid conformation closer to the channel’s central pore axis, thus priming the channel for activation. This idea is consistent with structural observations showing smaller relative motions of α1 versus β2 subunit M2-M3 linkers between closed and activated conformations of wild type α1β2γ2 receptors (Kim et al., 2020; Masiulis et al., 2019). There is also a trend towards enhanced activity upon introduction of proline at site 2 in the γ2 subunit, although the effect is much smaller than in α1. The weaker effect in γ2 could potentially reflect the expectation that the most probable receptor stoichiometry contains two α1 subunits and only one γ2 subunit (Chang et al., 1996; Laverty et al., 2019; Scott & Aricescu, 2019; Sigel & Steinmann, 2012). Substitution of the naturally occurring site 2 proline in the β2 subunit with serine also enhanced GABA-activation, but substitutions with alanine had no effect. Thus, specific sidechain interactions at this position in β2 are important, but proline is not a requirement. In summary, we show that absence of a proline at site 2 in the α1 subunit is important for channels to remain closed at rest. Our observations are consistent with the idea that the backbone geometry of the M2-M3 linker at site 2 in the α1 subunit is an important determinant of the channel’s activation energy landscape.

## Methods

### Mutagenesis and in vitro transcription

DNA for wild-type and mutant GABA_A_R rat α1, β2, and γ2 subunits was subcloned in the pUNIV vector (Venkatachalan et al., 2007). The mature protein numeration for β2 and γ2 subunits is the same for rat and human, but the numeration for the rat α1 subunit is (human numeration -1) for most of the subunit. Mutations were introduced using QuikChange II (Qiagen) or by GenScript and confirmed by sequencing of the entire subunit. Complementary RNA (cRNA) for each construct was generated (mMessage mMachine T7, Ambion), quantified (Qubit, ThermoFisher Scientific) and quality assessed (TapeStation, Agilent) prior to injection in *Xenopus laevis* oocytes.

### Isolation of *Xenopus laevis* oocytes and injection of cRNA

Defolliculated *Xenopus laevis* oocytes were obtained from EcoCyte Bioscience or harvested from mature female *Xenopus laevis* frogs (Nasco), which were housed in the University of Texas at Austin animal facility. Frog care and surgery followed the ARRIVE guidelines and the University of Texas at Austin IACUC-approved protocol. *Xenopus laevis* oocytes were harvested from frogs under tricaine anesthesia. A piece of ovary was removed from the frog, and placed in isolation media (108 mM NaCl, 2 mM KCl, 1 mM EDTA, 10 mM HEPES, pH = 7.5). Oocytes were manually isolated from the thecal and epithelial layers using forceps and then incubated in a collagenase buffer (0.5 mg/mL collagenase from *Clostridium histolytic*, 83 mM NaCl, 2 mM KCl, 1 mM MgCl_2_, 5 mM HEPES) to remove the follicular layer. Before injections, the defolliculated oocytes were transferred to a sterile incubation solution (88 mM NaCl, 1mM KCl, 2.4 mM NaHCO_3_, 19 mM HEPES, 0.82 mM MgSO_4_, 0.33 mM Ca(NO_3_)_2_, 0.91 mM CaCl_2_, 10,000 units/L penicillin, 50 mg/L gentamicin, 90 mg/L theophylline, and 220 mg/L sodium pyruvate, pH=7.5). Oocytes were injected with cRNA for α1, β2, and γ2 subunits (wild-type or mutants) in a 1:1:10 ratio (Nanoject, Drummond Scientific). Oocytes were incubated in the sterile incubation solution at 16 ºC.

### Two-electrode voltage clamp recordings

One to three days post injection, currents from channels expressed in *Xenopus laevis* oocytes were recorded in two-electrode voltage clamp (Oocyte Clamp OC-725C, Warner Instruments), digitized using a PowerLab 4/30 system (ADInstruments) and recorded using LabChart 8 software (ADInstruments). Data was obtained from at least two different batches of oocytes for each experimental group. Oocytes were held at -70 mV and perfused continuously (2 ml/min) with ND96 buffer (96 mM NaCl, 2 mM KCl, 1 mM CaCl_2_, 1 mM MgCl_2_, 5 mM HEPES, pH 7.5) or ND96 buffer containing picrotoxin (PTX), GABA, or GABA & propofol. PTX was diluted from a 0.5 M stock solution in DMSO. GABA was diluted from a 1 M stock solution in double distilled water stored at -80 °C. Propofol was diluted from a 30 mM stock solution in DMSO such that the final solution contained ≤0.1% V/V DMSO. The recording protocol was as follows: a 10 s pulse of 1mM PTX was followed by a series of 20-40 s pulses of increasing concentrations of GABA, a 20 s pulse of saturating GABA in combination with 30 µM propofol (PPF), and a final 10 s pulse of 1mM PTX. Every recording was bookended by applications of PTX to correct for any drift and to identify the zero current baseline. Pulses were sufficiently long to resolve the peak response and inter-pulse intervals were 5-15 minutes to allow washout with buffer and currents to return to baseline. Current traces were baselined by subtracting a spline fit to manually selected baseline regions in each trace. Baselined traces were then normalized by subtracting the zero current level in PTX and then dividing by the peak response in GABA and propofol. Baselining and normalization were done with custom scripts in MATLAB 2024b (MathWorks). Normalized GABA concentration-response curves (CRCs) were fit with the Hill equation:

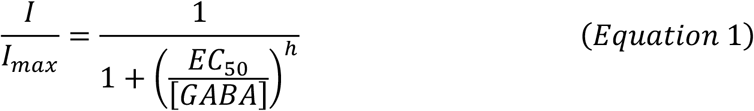

where *I* is the magnitude of the GABA-elicited current, *I*_*max*_ is the magnitude of the maximal current response elicited by the co-application of GABA and 30 µM propofol, [*GABA*] is the GABA concentration, *EC*_50_ is the concentration eliciting a half-maximal response, and *h* is the Hill slope.

### Statistical analysis

Summary data was analyzed using Prism 10 (GraphPad). Symbols and error bars are mean ± SEM, and box plots show median and interquartile intervals. Where applicable, we applied One-way Brown-Forsythe ANOVA followed by Dunnett’s T3 multiple comparisons test. Irrespective of *P*-value, we focus only on relatively large effects.

## Acknowledgements

This research was supported by NIH grant R01GM148591 to M.P.G-O. Additional support was provided by the University of Texas at Austin to N.G.D. We thank Dr. Susanne Ressl for helpful discussions during the conception of the study, and Anvita Bhatt for laboratory assistance.

## Author contributions

M.P.G.-O. conceived and supervised the project. N.G.D., P.G., and C.M.B. carried out, and N.G.D and C.M.B. analyzed, the two-electrode voltage clamp experiments. N.G.D and C.M.B produced the visualizations. N.G.D, C.M.B., and M.P.G.-O. wrote the manuscript.

